# Deep learning-based classification of resting-state fMRI independent-component analysis

**DOI:** 10.1101/2020.07.02.183772

**Authors:** Victor Nozais, Philippe Boutinaud, Violaine Verrecchia, Marie-Fateye Gueye, Pierre Yves Hervé, Christophe Tzourio, Bernard Mazoyer, Marc Joliot

**Affiliations:** Ginesislab, France; GIN, UMR5293, Bordeaux University, CNRS, CEA, France; Fealinx, Lyon, France; Bordeaux Population Health Research Center, UMR1219, Bordeaux University, Inserm, Bordeaux, France; Centre Hospitalier Universitaire de Bordeaux, France

**Keywords:** Resting-state, artificial intelligence, neuroimaging cohort, independent-component analysis, Brain functional network, classification

## Abstract

Functional connectivity analyses of fMRI data have shown that the activity of the brain at rest is spatially organized into resting-state networks (RSNs). RSNs appear as groups of anatomically distant but functionally tightly connected brain regions. Inter-RSN intrinsic connectivity analyses may provide an optimal spatial level of integration to analyze the variability of the functional connectome. Here we propose a deep learning approach to enable the automated classification of individual independent-component (IC) decompositions into a set of predefined RSNs. Two databases were used in this work, BIL&GIN and MRi-Share, with 427 and 1811 participants, respectively. We trained a multilayer perceptron (MLP) to classify each IC as one of 45 RSNs, using the IC classification of 282 participants in BIL&GIN for training and a 5-dimensional parameter grid search for hyperparameter optimization. It reached an accuracy of 92%. Predictions for the remaining individuals in BIL&GIN were tested against the original classification and demonstrated good spatial overlap between the cortical RSNs. As a first application, we created an RSN atlas based on MRi-Share. This atlas defined a brain parcellation in 29 RSNs covering 96% of the gray matter. Second, we proposed an individual-based analysis of the subdivision of the default-mode network into 4 networks. Minimal overlap between RSNs was found except in the angular gyrus and potentially in the *precuneus*. We thus provide the community with an individual IC classifier that can be used to analyze one dataset or to statistically compare different datasets for RSN spatial definitions.

## 1. Introduction

The resting-state is defined as a cognitive state of spontaneous activity that is not triggered by externally imposed tasks. In such a state, Biswal et al. (Biswal et al. 1995) used blood-oxygen-level dependent (BOLD) functional MRI (fMRI) techniques and showed for the first time that distant regions can have similar quasi-periodic low-frequency BOLD time courses. This interregional intrinsic connectivity phenomenon occurs between assemblies of regions that define so-called restingstate networks (RSNs). Interestingly, these networks are somehow related to the way the brain supports cognition, which was initially demonstrated by Smith et al. (Smith et al. 2009) by comparing the 30000-subject activation studies in the brain-mapping database (Laird et al. 2005) and a dataset of resting-state acquisitions. It has been demonstrated that intrinsic connectivity is a biomarker of events that occur throughout life, such as genetics (Richiardi et al. 2015; Kong et al. 2020), development and aging (Dosenbach et al. 2010; Zuo et al. 2010; Pervaiz et al. 2020), cognitive skills (Pervaiz et al. 2020) or mental content at the time of acquisition (G. Doucet et al. 2012). The signal-to-noise ratio of resting-state BOLD fMRI is very low and contaminated by numerous extraneural sources of noise. Following tailored postprocessing corrections, two ways of efficiently managing these problems have been identified: increase the number of participants and use spatial averaging. The former strategy is implemented in so-called cohort studies with thousands of subjects (Miller et al. 2016; Pervaiz et al. 2020), although this strategy is not always possible, especially when studying neural pathologies or rare types of brain organization. The latter strategy requires the definition of a parcellation of the brain gray-matter tissue in regions, networks, modules (G. Doucet et al. 2011) or systems (Fox et al. 2005). Based on our past work, we propose that the RSN level of integration can provide an ideal scale for describing the systemic brain organization of healthy individuals or patients.

To date, the definition of RSNs comes mainly from the analysis of subject groups that vary in number from tens to hundreds of subjects (M. P. van den Heuvel and Hulshoff Pol 2010; Yeo et al. 2011; G. E. Doucet et al. 2019; Pervaiz et al. 2020) primarily using group-based independent-component analysis (ICA) (Abou Elseoud et al. 2011; Beckmann et al. 2005; Calhoun et al. 2008; Damoiseaux et al. 2006; G. Doucet et al. 2011; Jutten and Herault 1991; Shirer et al. 2012; Smith et al. 2009) as well as other techniques (Chen et al. 2013; M. van den Heuvel et al. 2008; Varoquaux et al. 2011; Yeo et al. 2011). Under this framework, we previously proposed a methodology named MICCA (for multiscale individual component clustering algorithm (Naveau et al. 2012b)) to create an atlas based on individual subject ICA decomposition (Naveau et al. 2012a) using a hierarchical classification algorithm and ICASSO (Himberg et al. 2004). This methodology led us to propose an atlas with finer-grained partitions than were obtained from group-based processing methodologies. Note that this algorithm is not specific for ICA and can be applied to any technique that provides individual network-based decompositions of the data, e.g., seeding (Dosenbach et al. 2006; Margulies et al. 2007), snowball (Wig et al. 2014), restricted Boltzmann machine (Kim et al. 2020) or dictionary learning (Lv et al. 2015; Varoquaux et al. 2011).

In addition to the methodology used for their construction, the atlases are dependent on the population selected and the acquisition device. If some RSNs have been reproduced across many studies (M. P. van den Heuvel and Hulshoff Pol 2010), this reproducibility may be valid only up to some partitioning level and may strongly depend on the aforementioned variables. Using a badly fitting atlas can create strong biases in the analysis and lead to results that are difficult to interpret. The most straightforward example can be found in the analysis of patient or aged subject datasets. Because the majority of atlases are built from healthy young subjects, if one finds a decrease in the intrinsic connectivity between two networks in a patient compared with a healthy subject or in an older subject compared with young subjects, then this finding could be either interpreted as a true decrease or as a modification of the spatial support of one or both of the networks.

In addition to accommodating different populations, the price paid for fMRI sampling of the whole brain every few seconds (or even less) is dependent on the image geometry of the instrumentation, namely, the field homogeneity, antenna type, acquisition sequence, and head positioning in the antenna. Even if an atlas is used, we need a tool to adapt this atlas to the specificities of the scanner.

Atlases are very popular (for example, the regional anatomical-based atlas AAL (Tzourio-Mazoyer et al. 2002)). In addition to providing an anatomical reference, one of the main reasons for this popularity is that they fulfill the task of boosting the signal-to-noise ratio of both the task and resting-state fMRI by averaging the signal in each region. This benefit is potentially even higher with RSN atlases in which the number of voxels in each RSN is higher than in the regional atlases. Another benefit is that these atlases limit the number of tests that need to be calculated when using those variables for statistical analysis. The drawback is that the spatial topography of functional regions is strongly predictive of variation in behavior and lifestyle factors (Bijsterbosch et al. 2019; Bijsterbosch et al. 2018). The difficulty in interpreting variations of the intrinsic connectivity in term of spatial support or decreases (see above) implies that analyses based on individually defined RSNs may surpass atlas-based analyses and may possibly become mandatory in some cases (Bijsterbosch et al. 2019). Indeed, individual-based analyses provide support for new descriptions of the brain intrinsic connectivity organization, such as that proposed by Braga et al. (Braga and Buckner 2017) and Margulies et al. (Margulies 2017).

Individual-based decomposition can also be used to address the question of overlap, which indicates cases that show a region belonging to at least 2 RSNs. Most of the atlases of RSNs are built without allowing overlap. When calculating the intrinsic connectivity between 2 RSNs, overlaps are difficult to handle because they introduce some unwanted and nonbiological correlation between the 2 RSN BOLD signal variations. Moreover, these atlases do not acknowledge areas that could belong to two or more networks and represent potential hubs of connectivity when modeling the brain functional organization as a graph (Bullmore and Sporns 2009). Based on a group analysis, Yeo et al. (Yeo et al. 2014) and van den Heuvel et al. (M. P. van den Heuvel and Sporns 2013) demonstrated that there are in fact many regions of overlap between RSNs. The former group used a methodology where the constraint of unicity (no overlap) is probably less stringent than in the spatially independent ICA used by the latter, and they also performed a group analysis; however, assessing the region that truly belongs to 2 networks is difficult because of the overlap created by the group averaging. One way to solve this problem is to initially search the overlaps at the individual level and then perform a group statistical analysis.

Machine-learning algorithms, particularly artificial neural networks, have become more powerful in recent years, and applications in biological research have led to their successful application in medical imaging (Heinsfeld et al. 2017; Plis et al. 2014; Zhang et al. 2016). In this context, machine learning appears to be a potential tool to tackle the problem of automatic classification of individual structural or functional brain maps. Given a set of individual brain maps labeled in N classes, the goal is to train a supervised machine-learning algorithm to automatically classify each map of a new subject into one of the N classes. Some promising results in this area have already been obtained for a small number of classes with perceptrons ((Vergun et al. 2016) classification in 5 RSN classes) or convolutional neural networks (CNNs, (Chou et al. 2018; Lv et al. 2015; Zhao et al. 2018) classification in 10 RSN classes). Note that in those works, the ground truth used for training was given by a manual classification.

Our goal was to leverage the deep-learning methods to classify brain maps into a higher number of classes. In fact, the optimal number for ICA was proposed to be 45 RSNs (Abou Elseoud et al. 2011). Two datasets, namely, BIL&GIN (for Brain Imaging of Lateralization by the “Groupe d’Imagerie Neurofonctionelle”, 427 participants, (Mazoyer et al. 2016)) and MRi-Share (for Magnetic Resonance internet-based Student health research enterprise, 1811 participants, (Tsuchida et al. 2020)), obtained from two different populations and two different scanners, were used in this analysis. The ground truth was not provided by a manual classification but from the automatic clustering method of individual independent components (ICs) from 282 subjects of the BIL&GIN dataset using the MICCA (Naveau et al. 2012a; Naveau et al. 2012b) and ICCASO algorithms (Himberg et al. 2004). From this dataset, the number of classes was estimated at 45 (including some spatially reproducible artifacts). We first designed and trained a multilayer perceptron (MLP), a type of deep neural network (DNN), to automatically assign each IC extracted from individual fMRI sessions to one of the MICCA RSNs (or to a noise class), creating subject-specific functional brain maps without the need for manual labeling. Note that MICCA and MLP are complementary: the former is a clustering algorithm that cannot label new data, and the latter is a classification algorithm that cannot cluster the data by itself. In the second phase, the DNN was tested on the second part of the BIL&GIN dataset (145 subjects with unlabeled ICs), and the resulting labeling was tested against the MICCA labeling. In the third phase, the DNN predictor was applied to the MRi-Share dataset and once again compared with the MICCA labeling. As an application, this last analysis was used to provide both an atlas based on MRi-Share and an individual-based analysis of the subdivision of the default-mode network (DMN) of the brain. The DMN was chosen as it is specific to the resting state (Mazoyer et al. 2001; Raichle et al. 2001). In fact, its identification is overrepresented compared with the other networks in the ICA resting-state analysis. Based on an interindividual overlap analysis, we searched for regions that are part of 2 or more subsystems (Andrews-Hanna et al. 2010; Yeo et al. 2014) and could thus qualify as hubs (Bullmore and Sporns 2009).

## 2. Methods

Two databases, BIL&GIN (see 2.1) and MRi-Share (see 2.2), were used in this work (Table 1.). A graphical sketch of the analysis is presented in Figure 1, including the training of the MLP to select the best model (specific set of hyperparameters, see 2.4) using the MICCA-labeled ICs of BIL&GIN (see 2.3) and testing the MLP on 3 datasets (see 2.5).

**Fig. 1.**
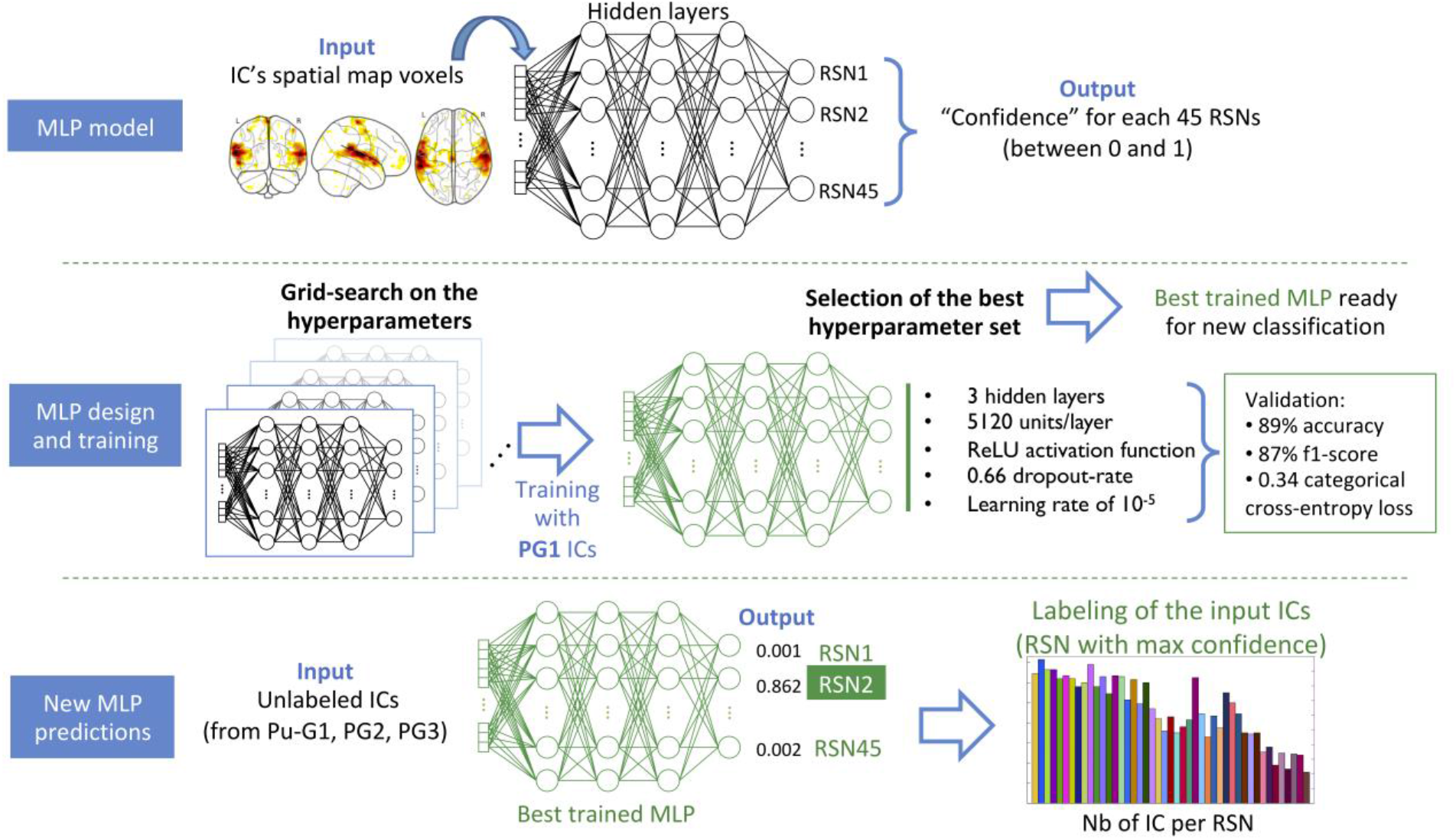
Top: Short description of the MLP principles. Middle: Grid search (testing all hyperparameter combinations to find the best one), which requires the training of each model using Ps-G1 ICs. The hyperparameters of the best MLP model and its evaluation metrics are given on the right side. This trained MLP model is the one used in subsequent classifications. Bottom: Using the trained MLP to classify unlabeled ICs and associate each IC with an RSN. On the far right, the number of PG2 ICs (Y-axis) classified in each RSN (X-axis) is shown as an example.

**Table 1.** Summary of the dataset. BIL&GIN: Brain Image Lateralization acquired by the “Groupe d’imagerie Neurofonctionelle”. MRi-Share: Magnetic Resonance internetbased Student health research enterprise. MICCA: Multiscale independent component clustering algorithm. IC: Independent component. G1/2/3: Group (of subjects) 1/2/3, respectively. Ps-G1 and Pu-G1: Independent components from G1 successfully and unsuccessfully labeled with MICCA, respectively. PG2/3: sets of Independent components extracted from groups 2/3, respectively.

| Subject groups<br>(num. subjects) | Dataset | IC groups for training and prediction |
| --- | --- | --- |
| G1<br>(282) | BIL&GIN | Ps-G1 (7999 ICs): subject ICs successfully labeled by MICCA, used for training<br>Pu-G1 (5730 ICs): subject ICs unsuccessfully labeled by MICCA, used in new predictions |
| G2<br>(145) | BIL&GIN | PG2 (7281 ICs): subject ICs unlabeled by MICCA, used in new predictions |
| G3<br>(1811) | MRi-Share | PG3 (90658 ICs): subject ICs unlabeled by MICCA, used in new predictions |

### 2.1. BIL&GIN dataset

The first dataset was from the BIL&GIN database (Mazoyer et al. 2016). This database was designed to investigate the cognitive, behavioral, and brain-morphological correlates of hemispheric specialization. A total of 427 participants who underwent both an anatomical MRI and a restingstate functional MRI were selected. Note that the BIL&GIN participant recruitment was biased toward young adults (age 27 ± 8 years [18-57], median of 24 years), balanced for sex (51% women, N = 219) and was enriched in left-handed individuals relative to the general population (46% versus 10%). The study was approved by the local ethics committee (Basse-Normandie, France).

Two independent data groups were defined: G1, 282 participants (age 26 ± 7 years [18-57]), and G2, 145 participants (age 29 ± 9 years [18-57]).

A full description of the BIL&GIN imaging dataset acquisition parameters and resting-state processing can be found in (G. Doucet et al. 2011; Mazoyer et al. 2016; Naveau et al. 2012b). A summary of the main parameters and steps of the analysis is given below.

#### Acquisition

Imaging was performed on a Philips Achieva 3-Tesla MRI scanner. The session started with acquisition of structural MR brain images, weighted volume (sequence parameters: repetition time (TR) = 20 ms; echo time (TE) = 4.6 ms; flip angle = 10°; inversion time = 800 ms; turbo field echo factor = 65; sense factor = 2; field of view = 256 x 256 x 180 mm^3^; 1 x 1 x 1 mm^3^ isotropic voxel size).

Spontaneous brain activity was monitored using BOLD fMRI, while the participants were at rest for 8 min T2*-echo planar imaging (sequence parameters: 240 volumes, 8 min; TR = 2 s; TE = 35 ms; flip angle = 80°; 31 axial slices; 3.75 x 3.75 x 3.75 mm^3^ isotropic voxel size). Immediately before fMRI scanning, participants were instructed to “keep their eyes closed, to relax, to refrain from moving, to stay awake, and to let their thoughts come and go”.

#### Processing

The analysis pipeline chains include temporal and motion correction of the fMRI signal and MNI stereotaxic normalization SPM5 (sampling 2×2×2 mm^3^) using the mediation of the T1-weighted anatomical device. Voxelwise temporal variance was removed using linear regression including the average white matter signal, average cerebrospinal fluid (CSF) signal and 6 movement parameters (3 rotations and 3 translations). The corrected signal was filtered temporally (0.01-0.1 Hz) and spatially (6 mm full width at half maximum (FWHM)). Individual fMRI data were individually processed using the ICA program called MELODIC (multivariate exploratory linear optimized decomposition into independent components, version 3.14) available in the FMRIB Software Library (FSL, (Smith et al. 2004)). The number of ICs was estimated by Laplace approximation (Minka and Thomas 2000). In the G1 subset, 13729 ICs were extracted (average of 49 ± 6 ICs per subject), and 7281 ICs were extracted in G2 (average of 50 ±7 per subject).

### 2.2. MRi-Share dataset

The second dataset was provided by the MRi-Share database (Tsuchida et al. 2020), a subpart of the i-Share (Internet-based Student Health Research Enterprise, http://www.i-share.fr), a large prospective cohort of French university students that investigates student health status (both physical and mental). MRi-Share was designed to investigate the brain morphological and functional organization of a subset of i-Share participants. We included 1,811 MRi-Share participants (referred to as G3) who completed full MRI examinations and did not show any abnormalities on their brain structural scans: the G3 group had an average age of 22.1 ± 2.3 years ([18-35], median at 21.7 years), a higher proportion of women (72%, 1,300 women) and 12.9% of left-handers (N = 233). The study was approved by the local ethics committee (Bordeaux, France).

#### Acquisition

Imaging was performed on a Siemens Prisma 3-Tesla MRI scanner. The session included structural MR brain images, including a high-resolution, three-dimensional MPRAGE T1-weighted volume (sequence parameters: TR = 2000 ms; TE = 2.03 ms; flip angle = 8°; inversion time = 880 ms; field of view = 256 x 256 x 192 mm^3^; 1 x 1 x 1 mm^3^ isotropic voxel size, in-plane acceleration = 2).

Spontaneous brain activity was monitored using BOLD fMRI while the participants were at rest to obtain 15 min multiband T2*-echo planar imaging (sequence parameters: 1058 volumes, 15 min; TR = 850 ms; TE = 35 ms; flip angle = 56°; 66 axial slices; 2.4×2.4×2.4 mm^3^ isotropic voxel size, x6 multislice acceleration (https://www.cmrr.umn.edu/multiband/).

Additional sequences, as part of the diffusion imaging acquisition protocol, were acquired to estimate a field map used for distortion correction (see below). Immediately before fMRI scanning, participants were instructed to “keep their eyes closed, to relax, to refrain from moving, to stay awake, and to let their thoughts come and go”.

#### Processing

The analysis pipeline concatenates distortion correction, motion correction and MNI stereotaxic SPM12 (https://www.fil.ion.ucl.ac.uk) normalization (sampling 2×2×2 mm^3^) using T1-weighted anatomical data. Voxelwise temporal variance was removed using linear regression, including the average white matter signal, average CSF signal, average gray matter signal, 6 movement parameters (3 rotations and 3 translations) and 18 parameters derived from the measured movement parameters (Friston et al. 1996). The corrected signal was filtered temporally (0.01-0.1 Hz) and spatially (5 mm full width at half maximum (FWHM)). Data were individually processed using the ICA MELODIC program (see above). A total of 90,658 ICs were extracted (average of 50 ± 5.5 ICs per subject).

#### 2.3. MICCA unsupervised labeling

Using an original unsupervised method named MICCA (Naveau et al. 2012b), RSNs were built using a spatial-overlap correlation criterion applied to the 282 individual ICA results from the G1 group. There are two outcomes of the MICCA algorithm: assignment of each IC to one of 45 classes plus one additional class and the creation of an atlas that describes the spatial support of these 45 RSNs (Naveau et al. 2012a), http://www.gin.cnrs.fr/en/tools/micca/). The additional 46th class, denoted “*class-0*”, includes all the ICs that could not be labeled with one of the 45 classes. Overall, 58% of the G1 individuals’ ICs were labeled and 42% remained unlabeled and were assigned to class-0. The former group of ICs, referred to as Ps-G1, consisted of 7,999 ICs, and the latter, Pu-G1, consisted of 5,730 ICs (“s” and “u” stand for successfully and unsuccessfully labeled by MICCA, respectively). This Pu-G1 class of unlabeled ICs was mostly composed of artifacts, but it also includes an unknown proportion of neural ICs.

Regarding the second outcome of the algorithm, the RSN atlas was built by creating for each RSN a voxelwise t-map of all ICs of Ps-G1 belonging to the same class. Figure 2 shows the cortical support of 29 of the 45 RSNs. Five other RSNs were localized in subcortical and medial temporal areas, and 2 were localized in the cerebellum. The 9 remaining RSNs were localized mostly in the white matter (1 RSN), in the temporal and frontal poles that are areas affected by susceptibility artifacts (2 RSNs) and in draining veins (4 RSNs). The remaining 2 RSNs showed band-like artifacts across both the white and the cortical matter and were identified as related to scanning defaults.

**Fig. 2.**
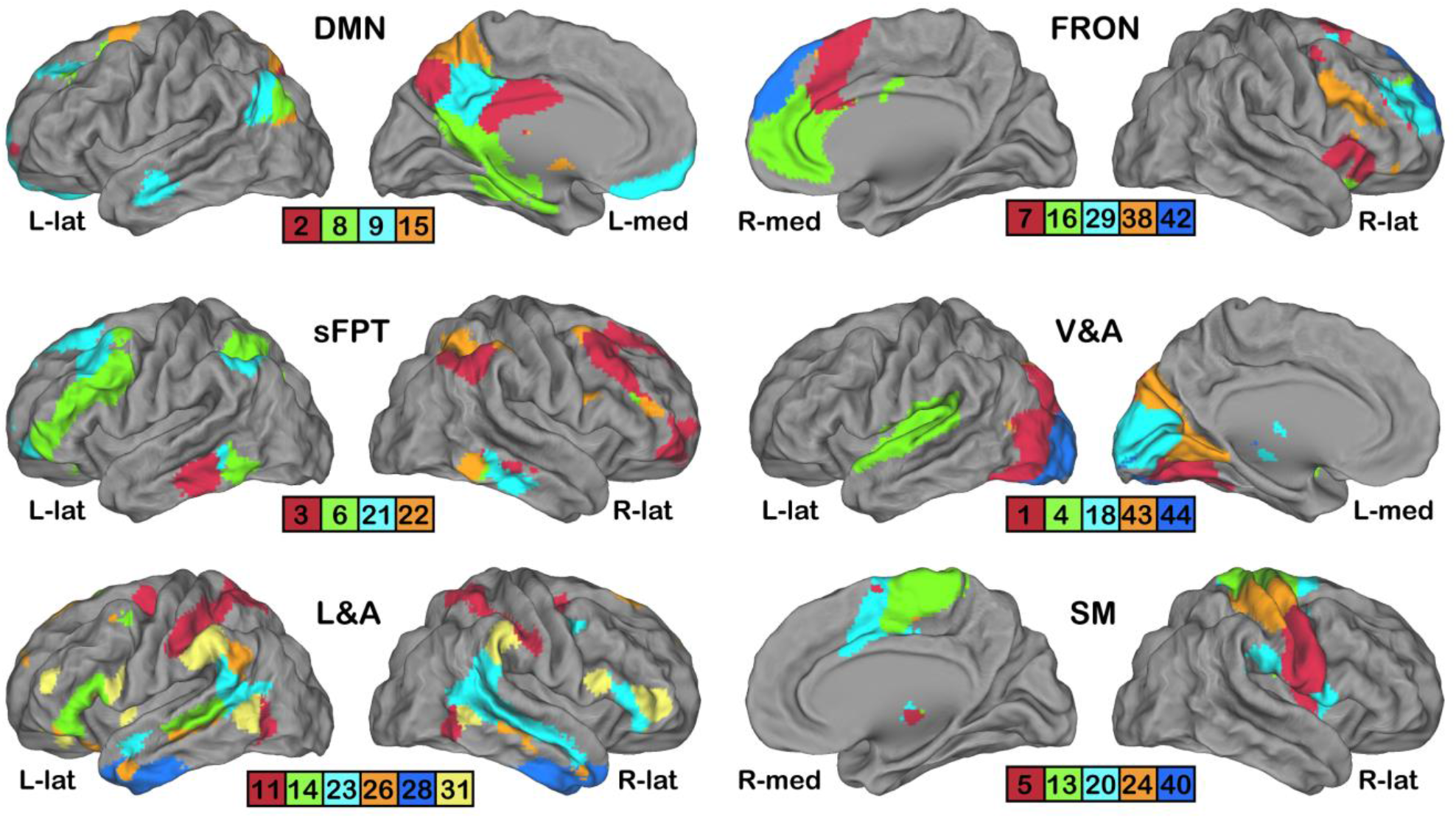
Twenty-nine color-coded cortical networks among the 45 RSN’s MICCA atlas. Each rendering set shows 4, 5 or 6 RSNs aggregated into 6partitions: DMN: default mode, sFPT: symmetric fronto-parieto-temporal, L&A: language and attentional, FRON: frontal, V&A: visual and auditory, and SM: sensorimotor. The renderings were computed using Caret software (Van Essen *et al*. 2001).

### 2.4. Artificial neural network specifications

Among machine-learning algorithms, two classes of DNNs are well adapted to multiclass monolabel classification problems such as ours: MLP and CNN. After testing different architectures (see the discussion for details), we selected the MLP approach. We implemented an MLP trained on the MICCA-labeled ICs (see 2.4.1.) using a 5-dimensional grid search (see 2.4.2). KNIME (Berthold et al. 2009) was used for the data management workflows; Python-based Keras (https://keras.io), Scikit-learn (Pedregosa et al. 2011) and TensorFlow (Abadi et al. 2016) were used for the DNN implementations; and Rstudio was used for visualization. All computations were run on a Centos computer with a Xeon ES2640 (DELL, USA), 40 cores, and 256 GB RAM and two NVIDIA P100 GPUs with 16 GB dedicated memory.

#### 2.4.1. Training and validation strategy

The Ps-G1 group of 7,999 ICs (see Figure 3A) was used as the training set of the DNN classifier. Each Ps-G1 IC consisted of a 3-dimensional z-map that was downsampled to the spatial resolution of the preprocessed fMRI data (~8 mm FWHM in each dimension) as 23×27×23 voxel-sized images. Note that the z-map represents a measure of the probability for each voxel belonging to this IC based on the normalized correlation between the voxel’s signal and the temporal IC signal extracted by the ICA (Beckmann et al. 2005).

**Fig. 3.**
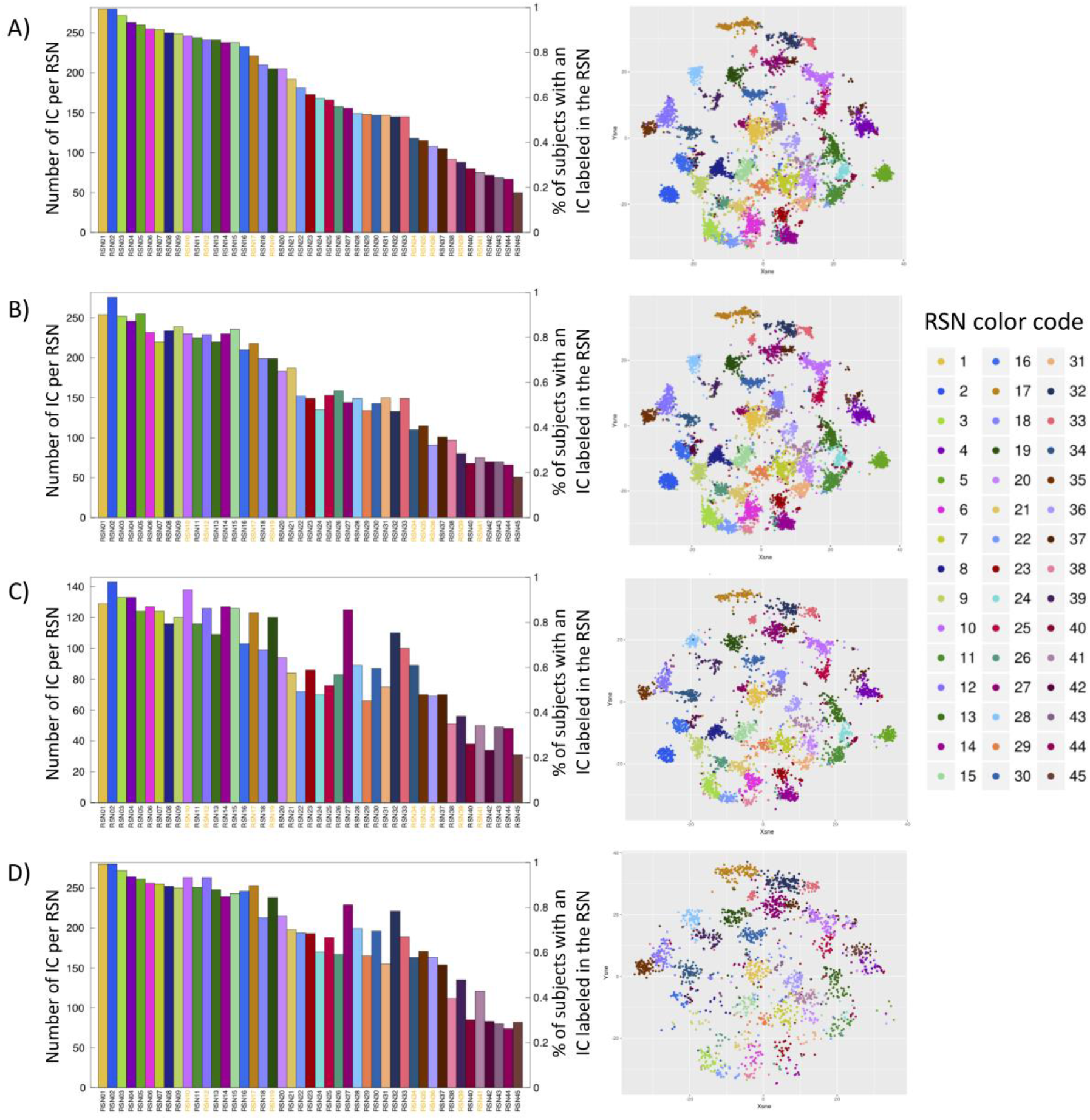
BIL&GIN processing. For each row, the figure on the left shows, for each RSN, the number of associated ICs (right Y-axis) and the equivalent % of labeled ICs per subject (left Y-axis); the figure on the right shows the t-SNE projection. A) MICCA labeling, B) DNN validation, C) PG2 labeling, and D) MICCA labeling and Pu-G1 combined. Only the Pu-G1 prediction was traced on the corresponding t-SNE graph. Note that the RSN names under the histograms were colored according to whether they were identified as a brain network (black) or an artifact (orange).

To quantify the accuracy of the classifiers, a stratified 5-fold cross-validation scheme was used. This method consists of dividing the data into 5 groups, or folds, of equal size. Each fold was selected in turn as a validation set, while the other 4 were used as training sets. Note that the ICs were balanced across RSN categories in the different folds, and the results showed that each fold validation set contained between 56 and 10 ICs for the most- and least-populated classes (RSN#01 and RSN#45), respectively.

#### 2.4.2. MLP model architecture and implementation

A grid-search approach was employed to search for the optimal values of 5 hyperparameters of the architecture of the MLP:

- Number of layers: [2; 3; 4]
- Number of units per layer (same for every layer): [512; 1,024; 2,048; 4,096; 5,120; 6,144]
- Type of activation function: [ReLU; tanh]
- Learning rate: [10^-5^; 10^-4^; 10^-3^]
- Dropout rate: [0; 0.25; 0.33; 0.4; 0.5; 0.66]

For each combination of hyperparameters (N = 648), a complete training and validation process was performed by the MLP. The metric used to assess the models’ quality was the average 5-fold validation loss. The hyperparameter combination giving the smallest loss was selected as the winning classifier.

The weights of the different hidden units were randomly initialized using a Glorot uniform initializer, which is also called the Xavier uniform initializer (Glorot and Bengio 2010). Categorical cross-entropy was used as the loss function, and Adam (Kingma and Ba 2014) was used as the optimizer. An adaptive learning-rate method was implemented that reduces the learning rate when the loss plateaus. To avoid overfitting the training data, two different methods were used. First, a dropout strategy was used, which consists of ignoring some randomly chosen units in each layer during the training. The ignored units changed in every iteration so that all of them were eventually trained. This method also helps the DNN learn more robust features (Srivastava et al. 2014). Second, an “early stopping” strategy was applied, which consisted of stopping the MLP training when the validation loss stops decreasing for a number of epochs, thus maintaining the loss at the minimum value and reducing overfitting. The trained model at the minimal validation loss epoch is then chosen. Additionally, because the classes showed an imbalanced distribution (Figure 3A), a class weight method was used, and it penalized errors more heavily for the underrepresented classes. The output dense layer uses the “softmax” activation function, with 45 units (one per class), to map the output as a confidence distribution for the predictions (mapped between 0 and 1). The final trained classifier is composed of 5 models, each trained on one of the 5 folds. For each RSN class, the confidence output by the classifier is the mean of the confidence yielded by those 5 models. The resulting predictions were used in the following analysis.

### 2.5. New classification and analyses

Once the model architecture of the MLP was defined and trained, we used it to classify 2 datasets that did not have previous RSN assignments.

The first dataset was composed of all ICs extracted from the BIL&GIN dataset that were not previously labeled by MICCA, namely, Pu-G1 (5,730 ICs) and PG2 (7,281 ICs). To test the robustness of the labeling, the spatial support of each RSN was compared to the spatial support of their MICCA-derived equivalent (see 2.5.1). Note that the Pu-G1 ICs were labeled to compare the labeling of all the ICs of G1 (Ps-G1 + Pu-G1) with PG2, as only part of the G1 ICs were labeled using MICCA (see 2.3).

The second dataset was composed of all ICs extracted from the MRi-Share database (PG3, 90,658 ICs). First, the spatial support of each RSN was compared to the MICCA RSN spatial support (see 2.5.1.), and then an RSN atlas was calculated (see 2.5.2). Finally, an individual-based analysis of the main subcomponents of the DMN subpartition was performed (see 2.5.3).

In all analyses following the MLP predictions, the ICs classified with a confidence below 50% were considered artifactual and reassigned to the noise-component class. Such a stringent criterion was also chosen to reduce the classification of ICs that overlap with only one part of the RSN. In fact, because of some instability in the automatic estimation of the number of ICs of the individual ICA decomposition, some RSN can be split into 2 or more ICs. Moreover, a unicity constraint was applied to the labeled data: for each set of ICs belonging to a specific individual, only one IC was assigned to each RSN. In cases where several ICs were competing for the same RSN, only the IC with the highest confidence was retained, and the others were discarded. This unicity method was used to make the results comparable to MICCA’s, since unicity was part of this algorithm.

#### 2.5.1. Spatial comparison of the MICCA and DNN results

We evaluated the differences in RSN neuroanatomical support depending on the algorithm used—MICCA or DNN—by quantifying the spatial overlap of PG2 MLP labeling (resp. PG3 MLP labeling) with G1 MICCA labeling. For this analysis, 2×2×2 mm^3^ sampling size images were used. To achieve this goal, a group-based voxelwise comparison was performed for each RSN using SPM12 (www.fil.ion.ucl.ac.uk/spm/), while balancing the samples in terms of the number of ICs, age, sex and handedness. Differences in overlap were computed and tested for significance (voxelwise t-test with p<0.05, familywise error (FWE) corrected).

#### 2.5.2. MRi-Share RSN atlas building

Based on the MRi-Share classification, an RSN atlas was created. First, all ICs (sampled at 2×2×2 mm^3^) belonging to the same class were averaged to define the spatial support of the corresponding RSN. For each RSN, the voxel distribution was fit by a mixture model (a Gaussian and two gamma functions as implemented in MELODIC (Beckmann et al. 2005)), and only the voxels above a 0.95 threshold were kept for further analysis. The atlas was built using a “winner take all” rule applied to each voxel, i.e., each voxel was associated with the RSN exhibiting the highest value.

#### 2.5.3. MRi-Share DNN-based default-mode network subpartitioning

Using the labeling results of the DNN classifier, the DMN subnetwork mapping was explored on the G3 dataset. An individual-based overlap analysis was performed instead of a group-based analysis to take into account overlaps at the individual level. We selected all RSNs that include part of the *precuneus*, which is considered to be the functional core of the DMN (Utevsky et al. 2014). Accordingly, we retained RSN #2, #8, #9 and #15 (see Figure 2, top row). Only the participants with an IC belonging to each of the 4 selected RSNs were considered in the analysis. Each individual IC (sampled at 2×2×2 mm^3^) was binarized using the associated mixture model and a threshold of 0.5, i.e., at the crossing between the positive gamma (modeling the active voxel) and the Gaussian distribution (modeling the noise). The mapping was performed by first associating each brain voxel, in each subject, to one of the 15 possible RSN “combinations”: none, 2, 8, 9, 15, 2&8, 2&9,…, 2&8&9&15. Then, the mapping itself consisted of selecting for each voxel the combination most often represented among the selected individuals. Note that voxels belonging to fewer than 50% of the individuals in each of the 4 RSNs were sorted into the “none” class.

## 3. Results

### 3.1. DNN training and validation

The grid search revealed several sets of hyperparameters with a comparable minimal loss (Figure 4A). The set with minimal loss was selected to define the best MLP: 3 layers of 5,120 units, with rectified linear unit (ReLU) activation with a 0.66 dropout rate and a learning rate of 10^-5^. This set achieved 89% accuracy and 0.34 categorical cross-entropy loss during crossvalidation (Figure 4A). Note that approximately half of the tested hyperparameter sets displayed a comparably low loss (0.36 or below), and the first 100 MLP models displayed a loss below 0.35 (see the enlargement of Figure 4A). This finding shows the remarkable stability of this type of algorithm when used for the type of classification problem encountered in studies such as ours.

**Fig. 4.**
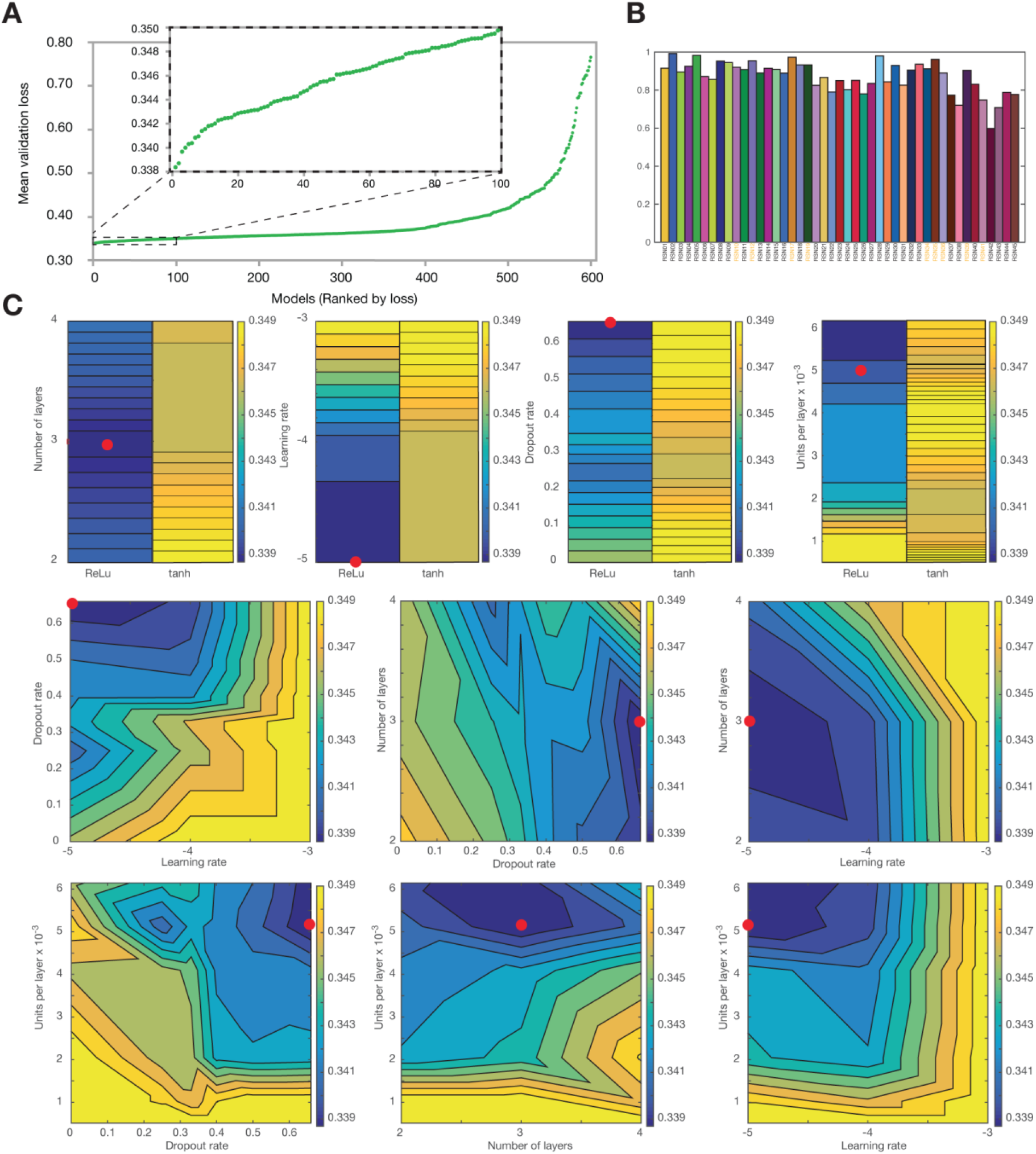
Analysis of the training set parameters. A) Sorted mean loss computed on each of the 648 MLP models tested in the grid search. Note that 48 models showing a loss greater 0.8 were not included in the figure. B) RSNf1-score of the selected model (minimal loss) across each RSN. Note that each RSN name is colored according to whether it is a brain network (black) or an artifact (orange). C) Interaction of the 5 hyperparameters on the loss; the losses were projected for each pair of hyperparameters. The red dots show the parameters of the optimal solution.

To observe the interaction of different hyperparameters on the loss, losses were projected for each pair of hyperparameters (Figure 4C), and all other hyperparameters were fixed at their value obtained from the best MLP. We observed that the activation function *ReLu* had better loss than *tanh*. The optimal number of layers was reproducibly found to be 3. Some of the hyperparameters (units per layer, dropout rate, learning rate) were chosen on the borders of the search space; outside the borders, technical limitations were met such as out of GPU memory, training not converging or maximum time set for hyperparameter optimization. As expected, there was a correlation between the number of units per layer and the level of the dropout rate: the more units per layer, the higher was the dropout rate needed to optimize the loss. Overall, we observed that the minimal loss was systematically located at the bottom of a well-defined “valley”.

To improve the model’s accuracy, we tested a consensus method aggregating the 5 best MLP models (according to the loss), which resulted in less than a 1% increase in accuracy. This minor improvement was achieved at the cost of drastically increasing the complexity and size of the training and prediction process; thus, the minimal-loss MLP was chosen for the testing analysis (see 3.2).

The selected MLP model showed a high and relatively stable f1-score across all RSNs (mean 0.87, SD 0.08; Figure 4B) despite a notable class imbalance (see Figure 3A). In addition, a 2-dimensional projection of the classification results with the t-SNE method (Maaten and Hinton 2008) shows a qualitative high similarity between the results obtained from MICCA (Figure 3A) and those from the validation steps of the MLP (Figure 3B). Note that the t-SNE coordinates were computed based on the matrix of all paired IC voxel value correlations from the full BIL&GIN dataset (Ps-G1, Pu-G1 and PG2). Additionally, the effects of applying the 0.5 confidence cutoff and the unicity constraint were tested on the validation data labeled with the MLP, since those post hoc steps were used in the rest of the study. This process led to a 1.5% increase in accuracy with the cutoff and an additional 1.7% increase in accuracy when applying both the cutoff and unicity, for a 3.2% increase in total, reaching an accuracy of over 92%.

### 3.2. Predictions on BIL&GIN

The selected optimal classifier was used to process both the Pu-G1 and PG2 BIL&GIN datasets. On PG2, 58% of the ICs were labeled to an RSN class (excluding Class-0). The *t-SNE* of PG2 (Figure 3C) was similar to both the *t-SNE* of MICCA labeling and that of DNN validation (Figure 3A & B). The percentage of detection per RSN of PG2 (Figure 3C) showed the same trend as that of MICCA (see Figure 3A) except for some networks, namely, RSN#27, #32, #33 and #34, which were overrepresented in the PG2 analysis. This phenomenon was expected considering that all the ICs of G2 were classified and only some were in G1. After adding the ICs labeled by MLP on the Pu-G1 ICs to the MICCA set, the overall percentage of labeled ICs in G1 reached 69%, and the detection percentage per RSN (Figure 3D) showed the same “overrepresentation” as in the G2 analysis.

The balanced group-based voxelwise comparison performed between PG2 MLP-labeled and Ps-G1 MICCA-labeled ICs did not show any significant voxelwise difference (p<0.05, FWE corrected) on any of the RSNs. Figure 5A shows the overlap analysis for 3 networks (RSN#09: default mode proper, RSN#11: dorsal attentional and RSN#14: language network). For the 45 RSNs, the average dice was 0.75 ± 0.01 (mean ± SD, N = 45), and when restricted to the 32 RSNs localized in the gray matter, it was 0.79 ± 0.08 (mean ± SD, N = 32). Note that among the latter RSNs, the lowest dice index was 0.50 for RSN#42, which primarily encompasses the frontal medial pole.

**Fig. 5.**
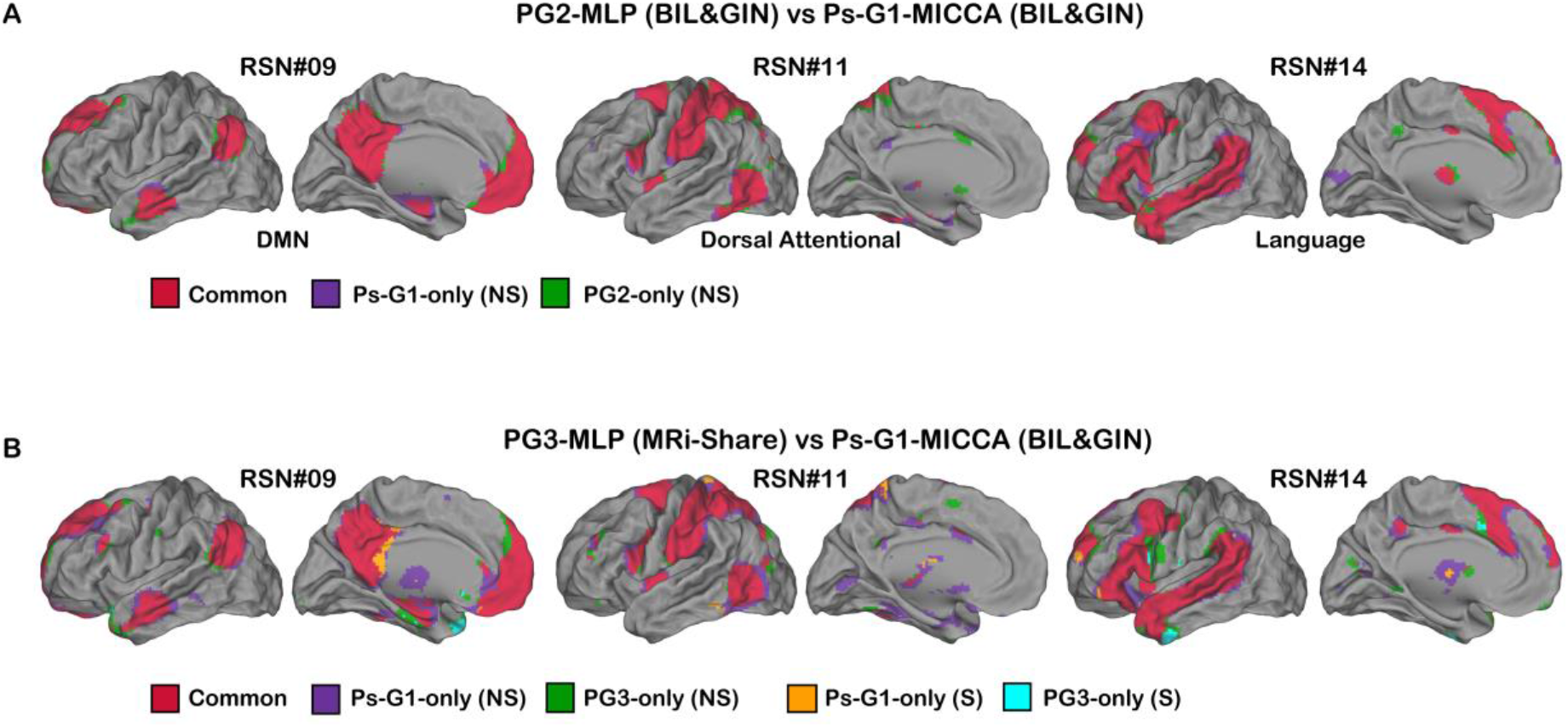
Balanced analysis of the overlap, based on a 0.05 FWE threshold, of the two dataset MLP classifications, i.e., PG2 (BIL&GIN, A) andPG3 (MRi-Share, B), with respect to theMICCA classification of Ps-G1 (named Ps-G1-MICCA). Red represents the common voxels, and purple and orange represent voxels with nonsignificant and significantly higher Ps-G1-MICCA values, respectively. Green and cyan represent voxels with nonsignificant and significantly lower Ps-G1-MICCA values, respectively. No significant difference was observed in the PG2-MLP vs Ps-G1-MICCA analysis. Note that only the left cerebral hemisphere is shown.

### 3.3. Prediction on MRi-Share (PG3)

#### 3.3.1. Spatial comparison of the MICCA and DNN results

The DNN classified 55.1% of the MRi-Share PG3 group ICs in one of the 45 RSNs (Figure 6, top). Among the 29 cortical networks of BIL&GIN, 27 RSNs appeared in the MRi-Share dataset analysis with a comparable or higher frequency. The 2 underrepresented networks were RSN#28 (localized mainly in the temporal poles) and RSN#42 (localized mainly in medial frontal areas). For the 27 networks, the balanced group-based voxelwise comparison between PG3 MLP-labeled and Ps-G1 MICCA-labeled results (see Figure 6, bottom) had an average dice index of 67.8 ± 4% (mean ± SD, N = 27). Figure 5B shows examples of the balanced group-based voxelwise comparison performed between PG3 MLP-labeled and Ps-G1 MICCA-labeled ICs for 3 RSNs (default mode, dorsal attentional and language networks). Compared with the same analysis with the PG2 BIL&GIN dataset (see 3.2), we only observed small significant differences between the spatial support of the RSNs. On all 27 cortical RSNs, the proportion of voxels appearing in MICCA but not in MRi-Share (p<0.05 FWE corrected) was on average 4.6 ± 3.1%, and the proportion of voxels in MRi-Share but not MICCA was 2.4 ± 2.6%. Those numbers defined on average an overlap percentage of 93 ± 4%. Among the subcortical (n=2) and temporal medial (n=3) RSNs, only RSN#25, encompassing the thalamus, appeared in MRi-Share with the same frequency as in MICCA. Regarding the other 11 networks, RSN#30, covering the cerebellum, appeared with a higher frequency in MRi-Share, while 9 networks, labeled as artifacts in BILGIN, exhibited a much lower frequency with a dice index below 50%, or were absent in MRi-Share.

**Fig. 6.**
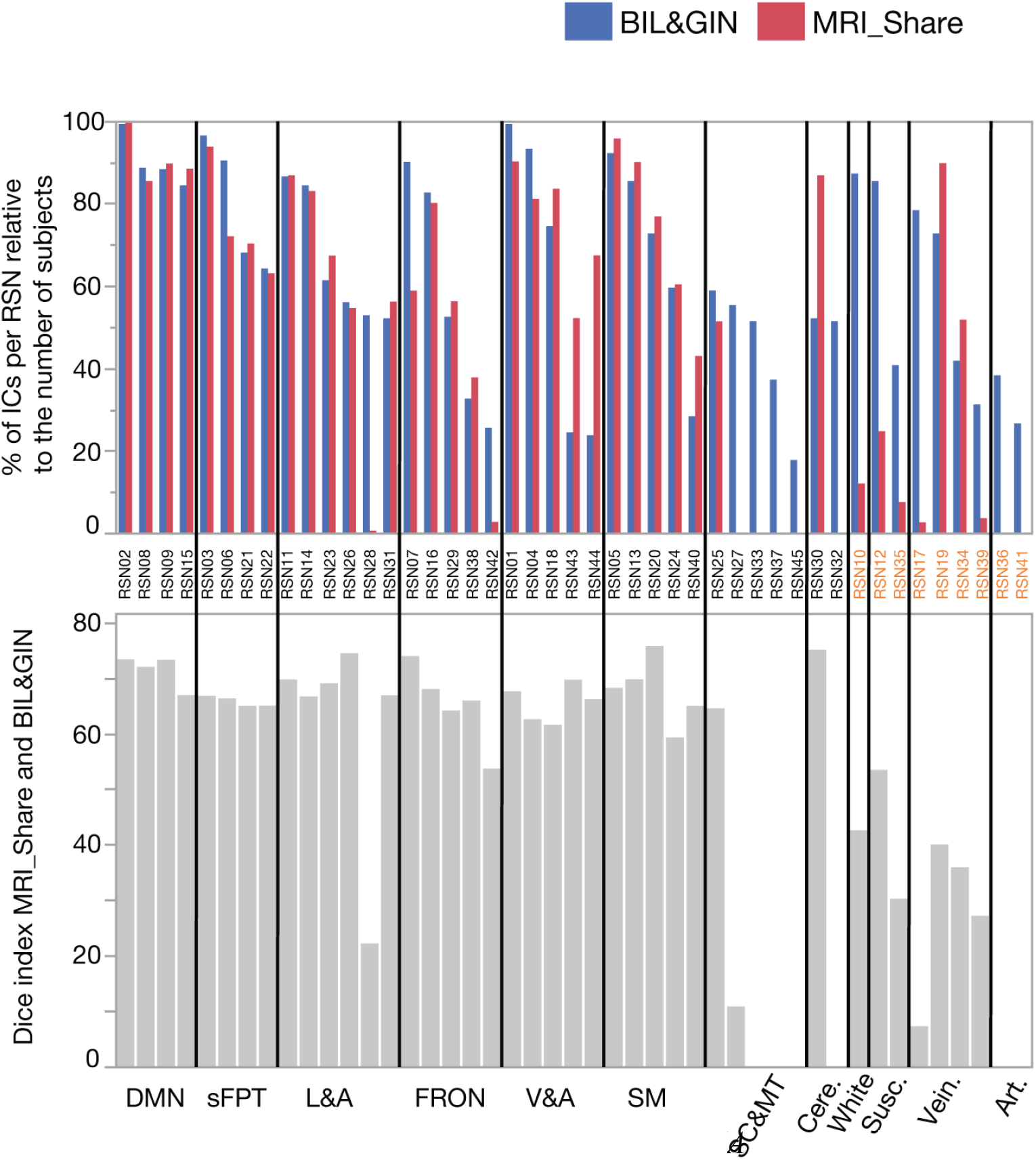
A) For each RSN, the % of labeled ICs per subject for MICCA (blue) and I-SHARE (red). B) Dice index of the overlap analysis of MICCA and MRi-Share based on the SPM balanced analysis results. The RSNs are ordered as in Figure 2 for the cortical networks: default mode (DMN, 4 RSNs), symmetric fronto-parieto-temporal (sFPT, 4 RSNs), language and attention (L&A, 5 RSNs), frontal (FRON, 5 RSNs), visual-auditory (V&A, 5 RSNs), and sensorimotor (SM, 5 RSNs). The other networks are in subcortical and medial temporal areas (SC&MT, 5 RSNs), in the cerebellum (Cere., 2 RSNs), in white matter (White, 1 RSN), or are susceptibility artifacts (Susc., 2 RSNs), draining vein (Vein., 4 RSNs) and scanner-related artifacts (Art., 2 RSNs). Note that each RSN name under the histograms was colored according to whether it was identified as a brain network (black) or an artifact (orange).

#### 3.3.2. MRi-Share RSN atlas

By using the MRi-Share PG3 MLP classification, a brain atlas (Figure 7) was built in the same manner as the original MICCA atlas (Figure 2). The atlas was built from the 27 cortical RSNs (Figure 7), with the addition of one RSN encompassing the thalami and one the cerebellum.

**Fig. 7.**
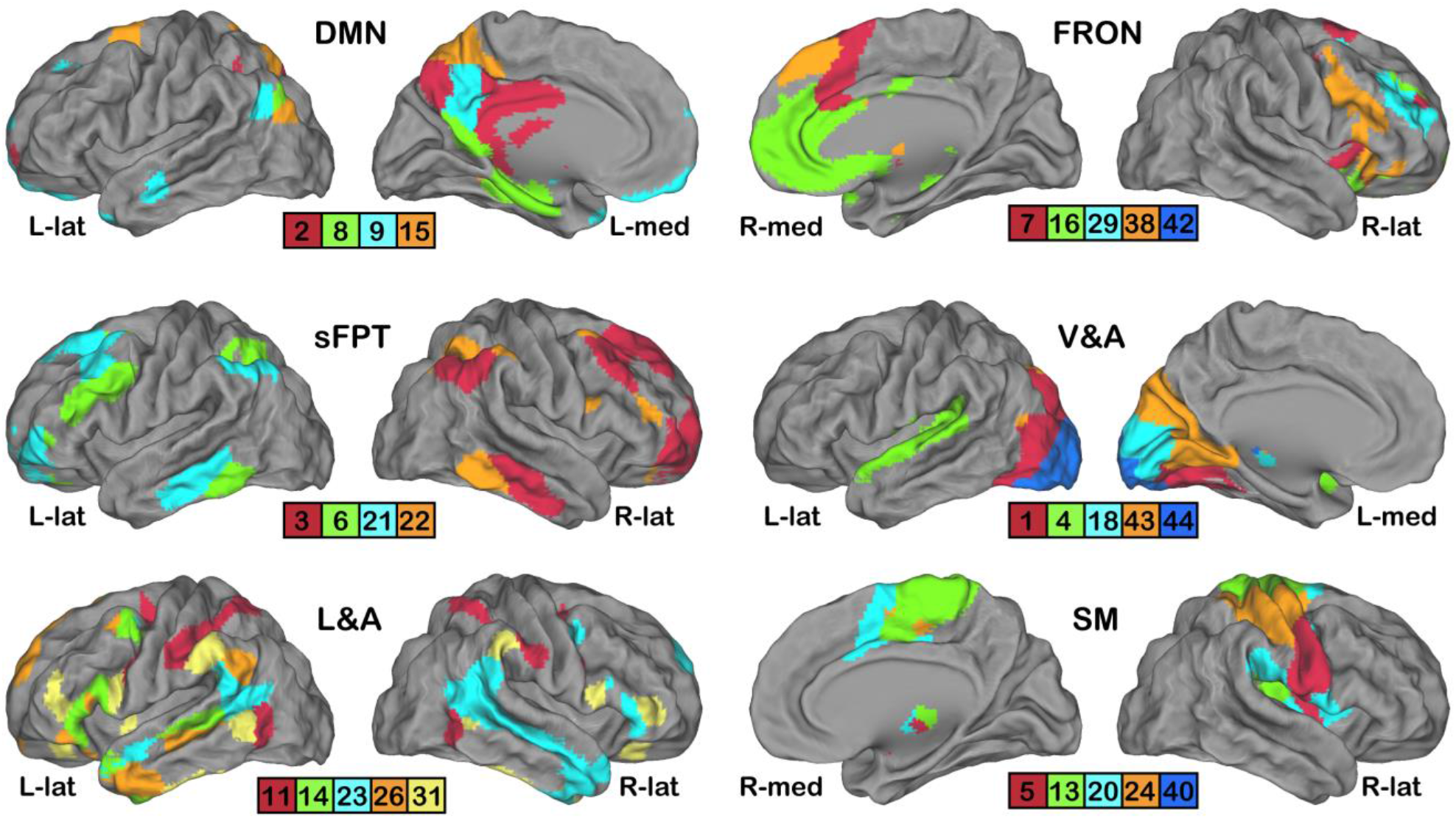
Twenty-seven cortical networks of the MRi-Share MICCA atlas. Each caret rendering set shows 4, 5 or 6 RSNs aggregated into 6partitions: DMN: default mode, sFPT: symmetric fronto-parieto-temporal, L&A: language and attentional, FRON: frontal, V&A: visual and auditory and SM: sensorimotor. For each partition and hemisphere, the most representative display (lateral or medial view) was chosen.

Note that the underrepresented RSNs (see 3.3.1.) were not considered because they were present in too few participants. Four percent of gray-matter voxels remained unlabeled; those voxels were mainly found at the lower part of the brain in areas exhibiting susceptibility artifacts. The main difference between the original MICCA and the MRi-Share-derived MLP atlas was first in the RSN#28 localized in the temporal poles. This network appeared in very few subjects in the MRi-Share dataset, and the temporal pole regions were affected to RSN#26 and RSN#23 in the left and right hemispheres, respectively. Similarly, the frontal-medial-temporal region of RSN#42 was taken over by another of the frontal networks (FRON), namely, RSN#38. The other difference in the symmetric fronto-parieto-temporal (sFPT) networks was the complete segregation between the left and right homotopic networks, while the temporal regions appeared on the contralateral hemisphere in BIL&GIN.

#### 3.3.3. Default-mode network subpartitioning

With 1,537 datasets showing the 4 DMN subnetworks (RSN#2, #8, #9, #15), G3 allowed us to accurately explore the overlaps between DMN subnetworks. The results showed a remarkably similar mapping between brain hemispheres (Figure 8) in two clusters, each showing all 4 RSNs: one centered on the *precuneus*, as expected, and another centered on the lateral parietal regions. The different networks were generally well defined with little overlap (Table 2) as ~82% of the voxels belonged to only one network. While overlaps of 3 (less than 1%) or 4 networks (0%) were minimal or null, overlaps of 2 networks (≅18%) were of 2 types. First, there were small bands of overlap at the border between 2 regions. These overlaps might have been created by the conjunction of spatial normalization and the spatial smoothing of the ICs (measured at FWHM = 6.4 mm in each orthogonal direction). The same might have occurred for the overlap between RSN#2 and RSN#9 in the left hemisphere, although it was less clear in the right hemisphere. Second, the overlap of RSN#8 and RSN#15 in lateral clusters centered on the left and right angular-2 regions (according to the AICHA atlas, (Joliot et al. 2015)) seemed to indicate that this region belonged to both networks.

**Fig. 8.**
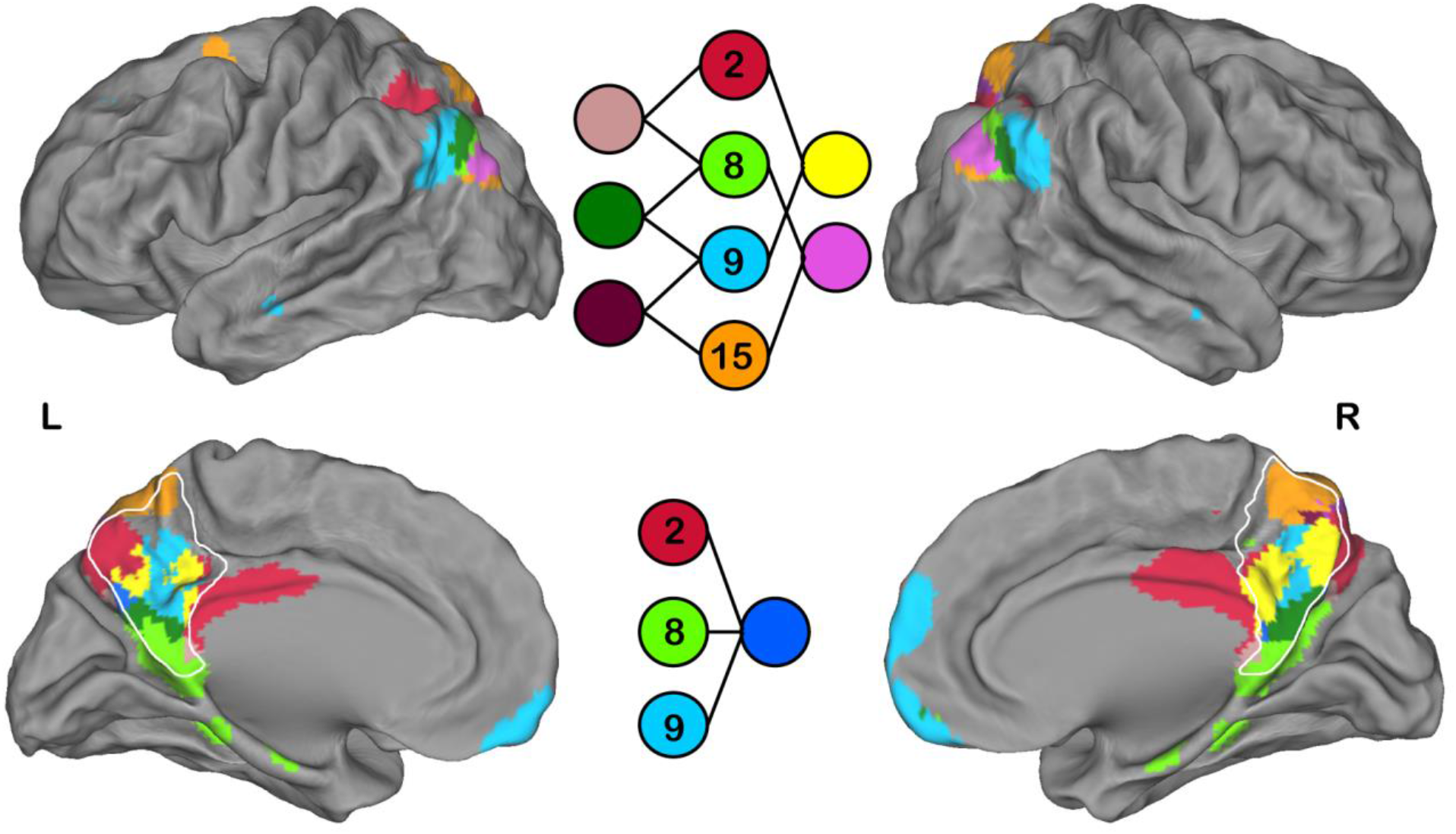
Overlap analysis of 4 DMN subnetworks identified with the MLP in the MRi-Share dataset. The main overlap can be found on the lateral surface RSN #8 and #15 (purple) and on the medial surface RSN #2 and #9 (yellow). Note that because of its small size, the overlap among RSN #2, #9, and #15 does not appear in the figure. The color legend is split into 2 parts to improve visibility. The white line delineates the precuneus.

**Table 2.** Overlap analysis of 4 DMNs in MRi-Share. From left to right: list of the 15 possible combinations (boldface), the absolute voxel count and the voxel count of each combination in terms of a percentage of the total voxel count. Dots indicate overlapping combinations without voxels.

| Combination<br>of<br>overlapping<br>RSN | Voxel count<br>(abs.) | Voxel count<br>(%) |
| --- | --- | --- |
| **R_2** | 3259 | 21 |
| **R_8** | 2541 | 16 |
| **R_9** | 3693 | 24 |
| **R_15** | 3250 | 21 |
| **R_2_8** | 290 | 2 |
| **R_2_9** | 868 | 6 |
| **R_2_15** | 253 | 2 |
| **R_8_9** | 540 | 3 |
| **R_8_15** | 835 | 5 |
| **R_9_15** | 55 | <1 |
| **R_2_8_9** | 73 | <1 |
| **R_2_8_15** | . | . |
| **R_2_9_15** | 1 | <1 |
| **R_8_9_15** | . | . |
| **R_2_8_9_15** | . | . |

## 4. Discussion

In this study, we explore the capacity of an artificial neural network to perform automatic classification of all individual independent components extracted by an ICA of resting-state fMRI data.

### 4.1. Choice of DNN methodology

As briefly explained in the section on the methods, two additional types of DNN were explored before deciding on the MLP model implemented in the present study. Both were CNN architectures, namely, VGG (Simonyan and Zisserman 2015) and ResneXt (Xie et al. 2017), which were updated to use 3D convolutions and batch normalization (Ioffe and Szegedy 2015). To determine which would be the most fitting, a scattershot approach was first used for hyperparameter selection on these three classical architectures.

The relatively small size of the dataset (training on 6,400 images, validation on 1,600 images) was more problematic for the CNN networks, which overfit quickly in fewer than 5 epochs. Moreover, this problem could not be overcome by traditional data augmentation methods, such as rotations, flips or translations. Indeed, the data here are represented as voxel high values in specific areas of the image, with all images normalized to the same MNI space during the preprocessing step. The different metrics used to evaluate the models also showed better results for the MLP compared with the CNN. For example, a nonoptimized MLP (ReLu activation function, 10^-3^ learning rate, 0.5 dropout rate, 2 layers and 512 units per layer) achieved a lower loss (0.38 vs. 0.52 for ResNeXt), a higher accuracy (88% vs. 85%) and a better f1-score (0.86 vs. 0.82). Those problems might have been resolved by fine-tuning the CNN hyperparameters; however, the CNN-based models took much longer to train than the MLP-based ones, and the grid-search approach applied to the MLP was not applicable to the CNNs in a sensible amount of time. In addition, the accuracy and loss results obtained with the first nonoptimized MLP were much more promising, which prompted our final choice.

In comparable classifications of individual-extracted RSNs, the CNN was used in two studies (Chou et al. 2018; Zhao et al. 2018). The reported accuracies were 99% for the average of 2 of the RSNs (Chou et al. 2018) and 95% for 10 RSNs (Zhao et al. 2018). On 45 RSNs, we reached an accuracy of 92%; however, when calculating the same index on the first 2 and on the first 10 most-populated classes, we achieved an accuracy of 95% and 93%, respectively. In another study, a perceptron-based classification in 5 RSNs (Vergun et al. 2016) showed a 90% accuracy (95% for us). Certain points make comparisons between these globally good results difficult. The 3 cited studies used manually labeled individual RSNs as the “ground truth”, while in our case, an automatic clustering method produced the ground truth. There is an advantage in the former methodology because only individual RSNs with a clear classification in one of the chosen RSN classes were used in the training and testing phases of the concurrent methods. In other words, certain classification mistakes likely occurred in our initial set of 7,999 individual ICs lowering the measured accuracy. In addition, because of the manual labeling, the number of classes was necessarily restricted and the classification problem was less complex than ours. Note that the proxy we used for reporting accuracy with a comparable number of RSNs (2/5/10) was still calculated on the classification of 45 RSN classes. For example, in the other studies, the DMN appears as one network, while in our cases, it appeared as 4 RSNs that share many spatial borders. Overall, we considered that our MLP-based method was in the same range of accuracy as the CNNs while handling at least 4 times more classes, and it was also superior to the previously tested perceptron.

In addition to the perceptron strategy, other machine-learning approaches have been tested by Vergun et al. (Vergun et al. 2016) on a limited dataset of 30 healthy individuals, including support vector machines (SVM), decision trees and Naive Bayes. Among those, the perceptron and SVM methods provided the best results (accuracy of 90%) but were only applied to a small number of classes (5). More generally, DNNs are better classifiers with large high-dimensionality data sets (Heinsfeld et al. 2017; Zhennan Yan et al. 2016).

### 4.2. Grid-search-based training

After selecting the DNN type, a careful choice of its hyperparameters was also needed. The gridsearch approach yielded several good options, among which the best was chosen (lowest mean loss on the validation folds). However, all the 100 best MLP models (i.e., MLPs with specific sets of hyperparameters) showed little difference in their mean loss compared with their variability, as illustrated by the difference between the maximal and minimal loss of each specific fold set (Supp. Figure 1). As a result, most of these models could probably be used interchangeably. The mean accuracy and mean f1-score of each model again showed a similar equivalence of the 100 best models, but they still indicated slightly better results in general for the best-ranked models. Overall, this finding demonstrated the high stability and accuracy of MLPs.

While almost all hyperparameters showed a clear and unique minimal loss, interestingly, the dropout rate systematically showed a local minimum between 0.2 and 0.3. Therefore, if speed is necessary or processing power is more limited, a dropout rate of 0.25 could be used (0.87 f1-score, equivalent to the chosen MLP model). Unexpectedly, the results thus showed that instead of the recommended 0.5 dropout rate (Srivastava et al. 2014), in our application, the best options were either higher or lower, highlighting the importance of the grid search (or equivalent procedure) in determining the optimal dropout rate. Additionally, the best learning rate and dropout rate were both located at extreme ends of the tested values, which indicated that even better results could have been obtained had the grid search been a little wider. However, further increasing the dropout rate or reducing the learning rate would lengthen the time needed to train the DNN.

We also tested an alternative hyperparameter optimization method, namely, sequential modelbased global optimization (or Bayesian optimization) (Bergstra et al. 2011), to determine whether the time needed to search for an optimal set of hyperparameters could be reduced. This random-search approach has recently been gaining popularity (Shahriari et al. 2016) and seems to be promising. Indeed, the grid search revealed that the hyperparameter space was mostly smooth, thus reducing the risk of the algorithm falling into a local minimum. Additionally, most of the hyperparameter space showed good results (more than 70% of the tested MLP models had a mean validation loss below 0.4), meaning that even if the Bayesian algorithm did not find the best set of hyperparameters, it should still be adequate for the present use. Using this optimization method, the optimum value was reached after testing only 120 models, compared to the 648 required in the traditional grid search. This dramatic reduction could therefore be used in similar applications in the future to increase the size of the searched space while reducing the time needed for the search and thus further optimize the DNN model used. Nonetheless, the grid-search approach was still necessary to control the validity of this new method. It also revealed the existence of a local minimum in the learning-rate space, which might be problematic in other projects (but not here) and should be kept in mind for future applications.

### 4.3. Prediction analyses

Once the DNN had been defined and refined, the first goal was to perform an array of tests to confirm its efficiency. These tests demonstrated the high f1-score of classification during training, equivalent t-SNE projections and quasi-perfect overlap mapping between data classified by MICCA and data from the same database (BIL&GIN) classified by the DNN. Moreover, its capacity to generalize from what it learned from MICCA-labeled ICs is a strength that becomes apparent with PuG1. This result showed that the DNN was capable of extracting relevant RSNs from the ICs discarded by MICCA, thus showing a higher classification performance, although most of those newly labeled PuG1 ICs were placed in categories that regrouped non-gray-matter areas. The only notable exception was RSN#27, which is located in the GM but in areas of susceptibility-induced artifacts. This finding makes it likely to be more variable than other RSNs, which leads to a lower detection rate with MICCA (because of the initial thresholding that excludes ICs with low spatial overlaps), while all the ICs were processed with the DNN.

Our second goal was to test our prediction DNN on another database acquired on a different scanner with a different MRI sequence and preprocessed by a different pipeline. The main differences between MRi-Share and BIL&GIN fMRI data acquisition are that the former uses a multiband sequence of a longer duration with better temporal and spatial sampling than the latter. During preprocessing, only the MRi-Share data were corrected for spatial distortions, and the spatial smoothing was ultimately weaker (FWHM of 6.4 mm, compared to 8.6 mm in BIL&GIN). Note that because of the multiband MRI sequence, the field of view for the MRi-Share data extended to the lower part of the head and encompassed the whole cerebellum, even in participants with very large heads. While the resultant classification and mapping were globally homogeneous for 27 of the 29 cortical networks in both datasets, two networks localized in the anterior medial frontal lobes, and the temporal poles (discussed below) were scarcely identified in MRi-Share. For the former, detection was low in BIL&GIN, and we do not have a clear explanation for this phenomenon except that the network is located in areas that were corrected for spatial distortions in MRi-Share but not in BIL&GIN. In addition to the abovementioned RSNs localized in the temporal pole, differences occurred in the lower part of the brain in 3 other RSNs. For those, further investigation will be required (see future works). On the one hand, the multiband accelerated sequence could have trouble reconstructing slices in areas affected by susceptibility variations. On the other hand, the networks covering both temporal poles were also associated with clusters in the outer CSF in BIL&GIN, thus making it a potential artifactual network (see 4.4 for further evidence). In addition to the 8 RSNs classified as artifacts, other missing or poorly overlapping networks were located inside the Sylvian fissure (2 RSNs) and the white matter (1 RSN). For the former group, this outcome was expected; some differences were probably related to the differences between the two MRI scanners that differentially affect the imaging of the vein and outer-brain artifacts located in the CSF. White-matter ICs were detected in a greater proportion in BIL&GIN than MRi-Share, which might be because the preprocessing was upgraded for the latter database to better remove signal variation in the white matter and ventricular CSF. This improvement in preprocessing could also explain why 2 RSNs located in the Sylvian fissures were not found. Even in BIL&GIN, they were already considered suspicious because they were found in a minimal number of participants and consisted of clusters centered on the intrasylvian CSF, but they were reported because of overlapping gray matter. Note also that in BIL&GIN, the spatial smoothness was higher than in MRi-Share; moreover, the subjects were young in both databases and thus did not exhibit aging-associated enlargement of the sulci. The filtering level difference was also responsible for a large proportion of the 5% of voxels observed in MICCA-BIL&GIN but not in DNN-MRi-Share, which presented voxels that mostly appeared as a thin band surrounding the area common to both analyses. Voxels found in DNN-MRi-Share but not in MICCA-BIL&GIN (3%) were aggregated as a small cluster in the lower part of the brain that could tentatively be ascribed to an increase in the signal-to-noise ratio because of the much shorter acquisition repetition time in MRi-Share compared with BIL&GIN (850 ms vs. 2000 ms).

### 4.4. Atlas creation

The first application led to the creation of a classical atlas, meaning without overlap between RSNs, using the 1,811 datasets of the MRi-Share database. The MICCA individual ICA-based methodology previously applied to the 282 subjects of the BIL&GIN database provided a parcellation in 45 RSNs, of which 29 were localized mainly in the cortical gray matter (Naveau et al. 2012a). In MRi-Share, the number of cortical RSNs decreased to 27. One of the “missing” network, strictly localized in the anterior frontal medial pole, was poorly detected in the MICCA labeling and was overlapping two other networks. Increasing the sampling size of the ICs could have solved this issue (see conclusion and perspective). The other missing RSN was localized in both the temporal poles and in the most frontal CSF in the BIL&GIN database. The MRi-Share dataset analysis suggested that this network was mainly related to susceptibility artifacts. Based on a winner takes all approach, the temporal poles were classified in MRi-Share as “language and attentional” networks, which makes sense since they included areas known to belong to the language network (Price 2012; Vigneau et al. 2006). This unmasking effect is a consequence of imposing the unicity for the voxel assignment to an RSN when creating an atlas. The unicity is used by all the atlases of RSNs (see review by (G. E. Doucet et al. 2019)) because it is complex to perform a statistical analysis on biomarkers calculated using overlapping networks and thus artifactually correlated signals. Compared with the literature, we found the 10 most-cited RSNs (see review of (M. P. van den Heuvel and Hulshoff Pol 2010)). However, due to the higher granularity in our study, each of those encompassed up to 4 of our RSNs. The DMN RSN overlapped 4 RSNs in our study, the primary motor RSN covered 4 of our RSNs, the left and right parieto-frontal RSNs each covered 2 of our RSNs and the 2 visual RSNs overlapped 4 of our RSNs. Among the additional networks, we found a network matching the location of the dorsal attentional network (RSN#11) that was surprisingly absent from the review by Van den Heuvel et al. (M. P. van den Heuvel and Hulshoff Pol 2010). Its hemispheric symmetrical localization may be the reason we were able to identify this network because other studies would split this network into the two fronto-parietal networks localized in each hemisphere. We also observed that those 10 networks did not cover the whole brain gray matter. With other algorithms, Yeo et al. (Yeo et al. 2011) and Gordon et al. (Gordon et al. 2015) covered most of the brain with 17 and 16 RSNs, respectively, and in both works, a match was found for the dorsal attentional network. With 27 identified cortical networks in our partition, the number of RSNs was higher than that chosen for those other works. However, according to Aboud et al. (Abou Elseoud et al. 2011), such numbers could be even higher than what they hypothesized, namely, an optimal group ICA model order of 80 RSNs leading to 45 gray matter RSNs. In our case, the number of classes was defined automatically but dependent on the number of ICs of the individual ICA (average of 50) that was computed with the Laplace approximation (Minka and Thomas 2000). Thus, this number of RSNs could probably be reached by manually setting the IC number in the ICA decomposition to a higher value. However, according to the tests performed by Abou Elseoud et al. (Abou Elseoud et al. 2011), it would also increase the risk of false positives. Note that one feature of our algorithm is that it allows for a subject to show only part of those networks because each IC is classified (or rejected) independently from the others. This behavior is reasonable when classifying individual ICA outputs because only part of the variability is taken in account in the decomposition (on average 40% in our case) or 2 or more RSNs can be aggregated in one RSN. Because of the classification, we can determine the potentially missing RSNs; a post hoc analysis could be designed to search for the missing RSNs. Of course, certain RSNs may not be functional and thus will be truly missing, or the spatial support may be different from that found in the general population.

To summarize, our algorithm provides the tools to study the individual variability of the brain restingstate organization. Even at the “atlas” level, we advocate that the improved robustness of mixing two databases is a strong incentive to further process other databases (see the future work section below).

### 4.5. DMN overlap

Overlap between RSNs has been scarcely studied in the literature except in a few studies, such as Yeo et al. (Yeo et al. 2014) and van den Heuvel et al. (M. P. van den Heuvel and Sporns 2013). Nevertheless, the implication of having or not having areas belonging to 2 or more networks is essential for understanding the resting-state brain organization, such as identifying these networks as a hub of IC in the graph analysis (Bullmore and Sporns 2009). Overlap analyses are strongly impacted by the number of RSNs, and in most atlases (see (G. E. Doucet et al. 2019) for a review), the DMN is not split into subcomponents. However, studies have demonstrated that in both animals (Margulies et al. 2009) and humans (Andrews-Hanna et al. 2010; Yeo et al. 2014), different parts of the *precuneus* belong to different networks that support different cognitive functions. In the study by Yeo et al. (Yeo et al. 2014), they described the DMN as split among 5 subnetworks and identified overlaps in the *precuneus*, the lateral temporal cortex, the posterior parietal cortex and the medial prefrontal cortex. Compared with their analysis, we found 2 overlaps at approximately the same location (*precuneus* and lateral parietal cortex); however, we were also able to more precisely identify their locations. In fact, they used a group analysis, which favors sensitivity over precision, while our individual-based methodology favors precision over sensitivity. In addition, their DMN definition does not completely match our definition. The van den Heuvel et al. study (M. P. van den Heuvel and Sporns 2013) uses both functional data (for extracting the 8-RSN decomposition) and diffusion imaging data (i.e., structural anatomy) for the definition of a hub, which is described as a region that links two or more RSNs. While showing the smoothness of the group analysis used to extract the RSNs, the hub of the *precuneus* showed a comparable localization to ours; moreover, we both found that the extension of the *precuneus* hub/overlap area was larger in the right than the left hemisphere. The lateral posterior parietal cortex also showed a greater overlap in the right than the left side, consistent with our findings and localization in the angular gyri. Consistent with Yeo et al. (Yeo et al. 2014), an overlap was also described in the medial prefrontal cortex, although it was not present in our results. This area was in RSN#16 in our study, and it mainly covers the anterior cingulate and has been linked by RSN intrinsic connectivity analysis to the salience networks but not to the DMN (see Figure 2, (G. Doucet et al. 2011)), and it does not extend to the *precuneus*. As shown in the supplementary material (Supp. Figure 2), its inclusion in the overlap analysis showed the same regional overlap in the medial frontal pole as that described in other studies.

Overall, in this overlap analysis of the subcomponent of the RSN, including the *precuneus*, it appears that using an independent-component analysis and individual overlap computation led to minimally overlapping ICs. In fact, while a group analysis showed a large overlap area extension, such an overlap was also observed in our study because of the spatial smoothing performed during data preprocessing to increase the signal-to-noise ratio.

## 5. Future work and conclusion

The present work has demonstrated the performance and robustness of the MLP for individual dataset classification. As such, it opens several lines of research that we intend to pursue. First, using recent GPUs with more memory, we will be able to apply deep learning methods on image data having better spatial resolution and thus to possibly perform a decomposition with a larger number of RSNs. Second, the atlas creation methodology will be upgraded to take into account the overlap analysis on the full set of RSNs. For this purpose, we hope to secure the authorization to use the largest available database, the UK-biobank (currently over 40K participants). To address the issue of networks covering mainly the lower part of the brain, which was identified in BIL&GIN but not in MRi-Share, we will also process databases with conventional functional sequences. We will also explore ways to increase the classification power, first by including the ICs discarded because of the unicity criteria, and then by analyzing the fraction of the variance that had been discarded in the first step of the independent component analysis. In summary, we provide the community with a new powerful tool for automatic classification of ICs, and although additional improvements are still planed, we demonstrate its utility in individual-based analyses.

## 6. Information Sharing Statement

The classifier and documentation are provided with the licensing Creative Commons Attribution-NonCommercial-NoDerivatives 4.0 International. Both BIL&GIN and MRi-Share data are open to collaborations and partnerships and supports local, national, and international collaborations from the public or private sector. For BIL&GIN requests to access the database should be sent through the GIN website (https://www.gin.cnrs.fr/en/current-research/axis2/bilgin-en/). For MRi-Share, requests to access the database should be sent at.

## 7. Acknowledgments

This work has been funded by a grant from “Agence Nationale de la Recherche” (Ginesilab, ANR-16-LCV2-0006), a University Bordeaux/CEA/CNRS and Fealinx joint laboratory. The i-Share cohort has been funded by a grant ANR-10-COHO-05-01 as part of the “Programme pour les Investissements d’Avenir”). Supplementary funding was received from the “Conseil Régional of Nouvelle-Aquitaine”, reference 4370420. The MRi-Share cohort has been supported by ANR-10-LABX-57 (TRAIL)

## 8. Ethics approval

The BIL&GIN study was approved by the ethics committee of Basse-Normandie (France). The MRi-Share study was approved by the ethics committee of Bordeaux (France).

