## Supplementary material for "Deep learning-based classification of resting-state fMRI independent-component analysis"

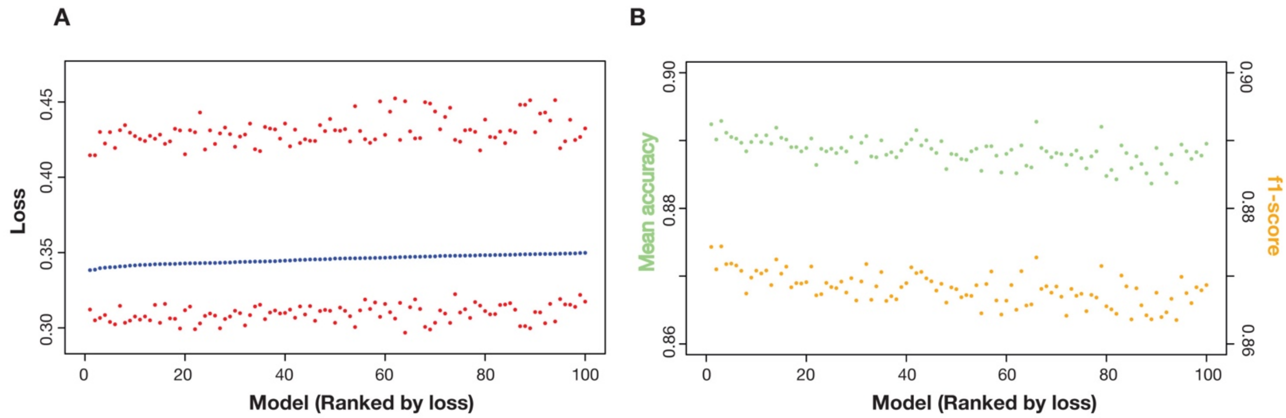


*Figure supp. 1: A) 100 best MLP models ranked by the average loss in the 5 folds. The red dots show the folds with the minimum and maximum loss. B) Accuracy and f1-scores computed on the validation set of the MLP solution ranked by the average loss in the 5 folds.*

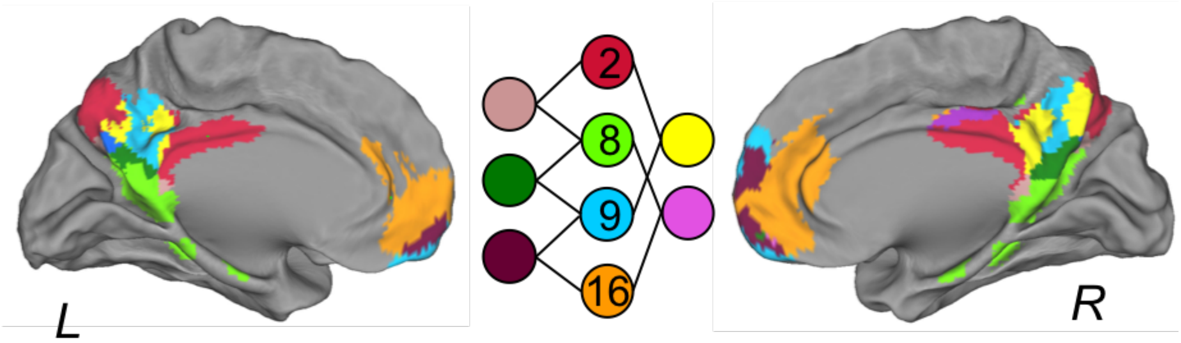


*Figure supp. 2: Overlap analysis of 3 DMN and one of the frontal medial (RSN#16) networks identified with the MLP in the MRi-Share dataset. Beside the already documented overlaps (see main text), we observed an overlap in a* medial frontal pole *overlap between the RSN#9 and RSN#16.*
